# A tale of two species: human and peafowl interactions in human dominated landscape influence each other’s behaviour

**DOI:** 10.1101/412254

**Authors:** Dhanashree Paranjpe, Priyanka Dange

## Abstract

Wild life is increasingly coming in contact with humans in many parts of the world. Human perception of wild life may be an important factor in population management and conservation practices in urban and semi-urban areas. Human attitude towards bird species may vary from non-violent coexistence to a perception of birds as pests. Based on the data collected from survey interviews, we studied the perceptions of local communities in Rajasthan, India towards Indian Peafowl. Local communities in Rajasthan reported Indian Peafowl as crop pest and this perception varied across seasons. The crop loss incurred due to Indian Peafowl also varied across seasons according to the respondents. Despite reporting peafowls as a crop pest, locals regularly offered grains for them as a traditional practice. Thus, at our study sites locals have mostly positive perception about Indian peafowl around them.

Food provisioning by local human population influenced diet composition and time-budget of Indian Peafowls at food provision and non-provision sites. Sites at which food provisioning was less or absent, peafowl spent more time in walking in search of food and more than half of their diet consisted of natural food. In contrast, the sites at which plenty of grains were available, time spent in walking was significantly less, while time spent in feeding was significantly more; and over 70% of their diet consisted of carbohydrate and protein-rich provisioned grains. Food provisioning changed the benefit: cost ratio (measured as time spent in feeding to time spent in all other behaviours) between provision and non-provision sites. Thus, food provisioning by humans can change feeding ecology of native Indian peafowl populations, while the presence of peafowl in human dominated landscape changes how humans interact with wild life around them.

## Introduction

Wild animals are increasingly finding themselves near human habitation due to scarcity of natural habitats, high human population density and changing land usage (Ayyappan et al. 2016; Pathak et al. 2002; Vijayan and Pati, 2001; Meena 2010). As a result, human-animal interactions and conflicts have risen. The impact of wild animals on local human population has mostly been studied in the context of human-wildlife conflicts (Agrawal et al. 2016) while, impact of local human population on animal populations living close to or within rural, semi-urban, and urban areas has rarely been studied.

There are potentially diverse types of interactions between humans and wildlife. The impact of human induced ecological stressors on animal populations in human dominated landscapes is becoming an important area of research (Sih et al. 2011; Agrawal et al. 2016). Wild animals can influence local human population socially, economically through crop loss, property damage, life loss, ecosystem services and aesthetic/ religious value. They can become sources of income through ecotourism, trade of animal derived products and so on.

Vicinity to human population can change natural behaviour and life history of animals in multiple ways. Human habitation or agricultural landscapes can serve as safer refuge areas or even new habitats for wildlife (Horgan et al. 2017). Humans often provide food to wildlife voluntarily around their homes, in religious places as a ritual or in wild life tourism spots/ feeding stations (Robb et al. 2008; Sengupta et al. 2015). In addition to this, indirect food provisioning in the form of crops, ornamental plants, livestock or waste food thrown in open becomes easy and reliable source of food for wild life. Food provided by humans tends to be calorie rich, easily digestible, available at predictable times and places. It is known that food provisioning might change feeding habits, diet preference or diet composition of wild mammals (Robb et al. 2008; Sih et al. 2011; Sengupta et al. 2015; Ayyappan et al. 2016; Sengupta and Radhakrishna 2018). Reduced predatory pressure and regular availability of nutrient rich food (in the form of crops) round the year are likely to provide greater resilience of wild animals. It also allows them to live successfully close to agricultural landscape and human habitation (Gilroy and Sutherland 2007; Sih et al. 2011; Ayyappan et al. 2016).

There are many avian species like crows, sparrows, pigeons, cranes, egrets that inhabit human-dominated landscapes or stay very close to / within human habitation. Appreciation for birds and even individual species may vary from nonviolent coexistence (including feeding them) to a perception of birds as pests (Clergeau et al. 2001). Very little work has been done on how human perception/ attitude towards avian species can influence the outcome of their interactions with humans. A multi-dimensional approach to study such interactions is necessary in order to understand the capacity of local stakeholders to accept wild species (Carpenter et al., 2000) as well as the potential effect of these interactions on both humans and wildlife who share common spaces.

Indian peafowl (*Pavo cristatus*) is a species that has co-habited human dominated landscapes for centuries in its native geographical range. This avian species is native to Indian subcontinent and has been introduced in many parts of the world relatively recently. Although, it’s native habitat is undergrowth in open forest and woodlands near a waterbody, it is also known to occur near farmlands, villages and increasingly becoming common in urban and semi-urban areas (Burton and Burton 2002). In some places, such as Morachi Chincholi in Maharashtra, India, “eco-tourism” based on Indian peafowl is flourishing and it is an additional or alternative source of income for local community (personal observations). In contrast to this “positive” impact, anecdotal reports suggest (Ganesan, R. 2016) that the Indian peafowl can cause substantial crop losses in the areas where their population density is high. There are no known studies from India that estimate the crop losses due to peafowls. It is not known whether peafowl really affect crops like other known crop pests (e.g. wild boar, elephants, deer, etc.). Numerous reports can be found in newspapers in India about deaths of individuals or groups of peacocks. The deaths may be due to natural reasons such as predation, water scarcity or may be the result of unintentional poisoning due to insecticides/ pesticides sprayed on crops that the peafowl later feed on. Sometimes intentional killing of Indian peafowl by certain tribal communities for their ornamental feathers and meat has also been suspected/ reported. In contrast to this, in some parts of India (e.g. State of Rajasthan), peafowl are believed to be sacred and people actively offer grains for them as part of their daily ritual. These varied interactions between Indian peafowl and local human population make it an interesting study system to understand consequences of human-wild life interactions. It remains to be seen how common are the positive perceptions and associated beliefs/ rituals throughout the region in which the population density of Indian peafowl is higher. Can these perceptions result in curbing population decline or effective management/ conservation of the species? To address these questions we studied three aspects of human-peafowl interactions in details. The objectives of this study were: (i) To understand the perceptions of local community about Indian peafowl, (ii) To estimate impact of Indian peafowl populations on local agriculture and (iii) To estimate the impact of food provisioning by local community on peafowl populations.

## Materials and methods

### Study area

We selected three field sites-Morachi Chincholi, Nashik and Rajasthan (Table 1). Selection of study areas was based on several criteria such as their closeness to human habitation, accessibility throughout the year, history of peafowl populations in the area, potential interactions between humans and peafowl (food provisioning, peafowl visiting/ roosting on agricultural land and village homes, sell of peafowl feathers or related “products”, tourism, cultural significance), etc. Areas located outside protected areas and close to human habitation were given preference.

**Table 1:**
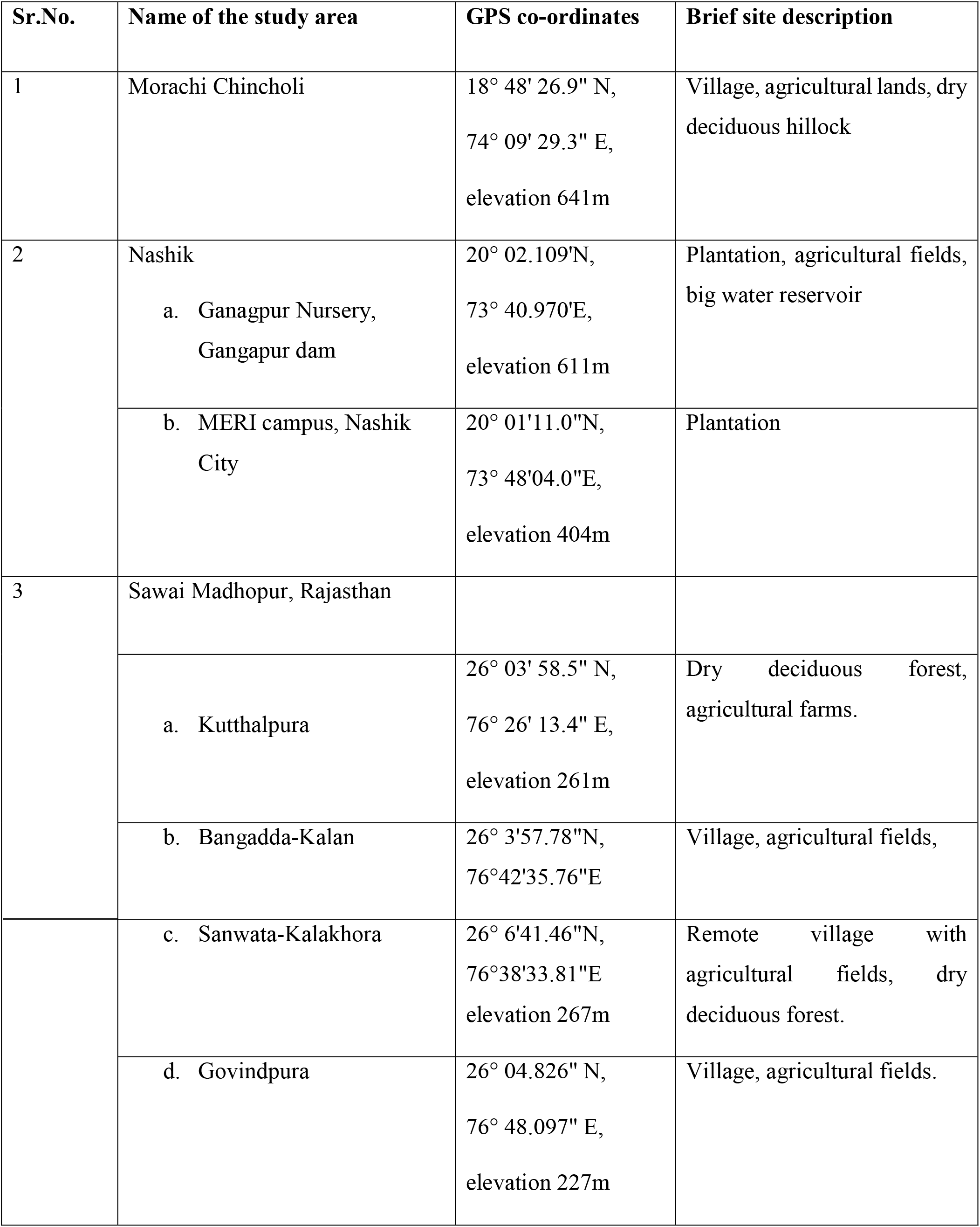
Brief description of study areas

We used questionnaire based interviews (N = 196) to estimate impact of Indian peafowl populations on local agriculture and to understand perceptions of local people about Indian peafowl in four villages located on the periphery of Ranthambhore Tiger Reserve (RTR), in the state of Rajasthan, India. Before actual survey, a pilot survey was conducted to standardize the protocol and contents of the questionnaire. Based on the results of the pilot survey some questions were modified or added to reduce ambiguity in questions and get more accurate data. Interviews were conducted by taking family as a unit respondent. The questionnaire included fixed and open-ended questions (See supplementary material for complete questionnaire). Interviews started with the interviewer clearly explaining objectives of the study and the interview was continued only if the respondent(s) gave consent. The questionnaire consisted of 3 parts: (i) basic information about the respondent such as name, age, address, number of people in family unit, source(s) of income (agriculture/ others). (ii) Questions related to perceived impact of peafowls on agriculture (iii) questions related to social/ cultural/ traditional perceptions about Indian peafowl. Interviews were conducted in three different seasons: November 2016 (harvest time for kharif crops, winter season, N = 55), March-April 2017 (Harvest time for rabbi crops, beginning of dry summer season, N = 86), July-August 2017 (standing crops, monsoon season, N = 55) to check if responses to some of the questions related to agriculture differed according to stage of crop/ season. For each survey equal number of respondents were sampled randomly from the local village populations.

To estimate the impact of food provisioning by local community on peafowl populations, behaviours of the Indian peafowl were studied at Morachi Chincholi, Nashik and Rajasthan throughout the year using direct field observations. Observations were recorded during morning (6 am to 10 am) and evening (4 pm to 7 pm). Behaviours were also observed in the late-afternoons if peafowl were found to be active. The behavioural observations of peafowl were noted while patrolling the study area using continuous scan and focal animal sampling. Focal animals were chosen randomly from a group (if the peafowl were seen in a group). Additionally, opportunistic observations were recorded for individuals sighted during the walk around the study sites. Behaviours of focal animal were recorded continuously without disturbing or going too close to focal animal (typically from a distance of 10-100m) until they were out of the sight. During feeding bouts, location of feeding was recorded and was later classified as provisioned and non-provisioned sites. Gender of the focal animal was noted along with the date, place and the total duration of observations. Data were also collected about the type of food the peafowl ate during the observations using binoculars to get clear view of the type of food they were eating. Observations were stopped when the focal animal flew/ walked out of observer’s sight.

For calculating time budget of the Indian peafowl populations, the observer(s) documented start/ end/ transition of behaviours of focal animal as a running commentary using handheld digital audio recorder (SONY ICD-UX560F). Occasionally, data were recorded as videos using a Panasonic camcorder (Full HD, 29.8mm wide 20.4 megapixel HC-X920) if a large group of peafowl was found near feeding site.

### Data processing

The types of food items peafowls ate were listed using direct observations (Table 2). Cereals, pulses, vegetables and processed food is available to peafowl only around human habitation and hence was considered as provisioned food, while all other food items such as grass, leaves, sprouts, fruits, seeds, insects, worms were considered as natural food items (Table 2). Based on direct observations of peafowl eating various food items, proportion of times it was seen eating natural and provisioned food items in peafowl diet was calculated. If it was not clear what the peafowl were eating, such observations were not considered for analysis of diet composition. The food items were further categorised as cereals (grains such as sorghum, wheat, bajra, rice, corn, etc.), pulses (split pigeon peas, black gram lentils, black-eyed pea, Indian brown lentils, green gram, peanuts, etc.), non-grains (fruits, grass, seeds, leaves, insects, vegetables, etc.) and processed food. Number of times peafowls were seen eating each food type was counted. Based on this frequency, the percentage of each food type in their diet was calculated.

**Table 2:** Diet composition (presence/ absence of provisioned and non-provisioned food) of Indian Peafowl *Pavo cristatus* at three field sites based on direct field observations.

| Diet Item | Food Category | Natural/<br>provisioned<br>food | Morachi<br>Chincholi | Nashik | Rajasthan |
| --- | --- | --- | --- | --- | --- |
| Sorghum (*Sorghum spp*) | Cereals | P | 1 | 0 | 0 |
| Wheat (*Triticum spp.*) | Cereals | P | 1 | 1 | 1 |
| Bajra (*Sorghum spp*) | Cereals | P | 1 | 0 | 1 |
| Rice (*Oryza sativum*) | Cereals | P | 1 | 1 | 1 |
| Corn (*Zea mays*) | Cereals | P | 1 | 0 | 0 |
| Unknown grains | Cereals | P | 1 | 0 | 0 |
| Vilayati Chinch<br>(*Pithecellobium dulce* ) | Non-grain | N | 0 | 1 | 0 |
| Methi Grass | Non-grain | N | 1 | 0 | 0 |
| Insects | Non-grain | N | 1 | 1 | 1 |
| Leaves/ grass | Non-grain | N | 1 | 1 | 1 |
| Seeds | Non-grain | N | 0 | 1 | 0 |
| Chillies (*Capsicum*<br>*annum*) | Non-grain | P | 1 | 0 | 1 |
| Vegetables | Non-grain | P | 1 | 0 | 0 |
| Unknown (non-grain) | Non-grain | N | 1 | 1 | 1 |
| Processed food | Processed food | P | 1 | 1 | 0 |
| Indian Brown lentils | Pulses | P | 1 | 0 | 1 |
| Green gram<br>(*Vigna radiata*) | Pulses | P | 1 | 0 | 1 |
| Split pigeon peas<br>(*Cajanus cajan*) | Pulses | P | 0 | 0 | 1 |
| Black gram lentils<br>(*Vigna mungo*) | Pulses | P | 1 | 0 | 0 |
| Moth beans<br>(*Vigna aconitifolia*) | Pulses | P | 1 | 0 | 0 |
| Black eyed peas<br>(*Vigna unguiculata*) | Pulses | P | 1 | 0 | 0 |
| Chickpeas<br>(*Cicer arietum*) | Pulses | P | 1 | 0 | 0 |
| Peanuts | Pulses | P | 1 | 0 | 0 |

The ethogram of the peafowl comprised of following behaviours: alert, walk, peck, preen, jump, run, scratch, calls, shake, scrape, search, turn, flap, fly, stand, display related, climb, headshake, perch, chase away, sit, pause, poop, drink, aggressive and shiver. (Fig.1). Relative frequency of each behaviour recorded for 293 individuals was calculated as (# of times a behaviour is shown/total no. of observations)*100 to identify the major and minor behaviours of Indian Peafowl from the ethogram. Peck, Alert and Walk were the major behaviours while the minor (<10% frequency) were pooled as other behaviours. For calculating time budget of the Indian peafowl populations, data about time of beginning of a behaviour and end of behaviour/ transition to other behavioural state was noted down from the audio records. This was used to calculate the time spent in various activities/ behaviours for each recorded individual. Fraction of time spent in each activity was summed across all individuals in the population. Time budget of the whole population was calculated as time spent in each activity divided by total time of observation for the population.

**Fig1.**
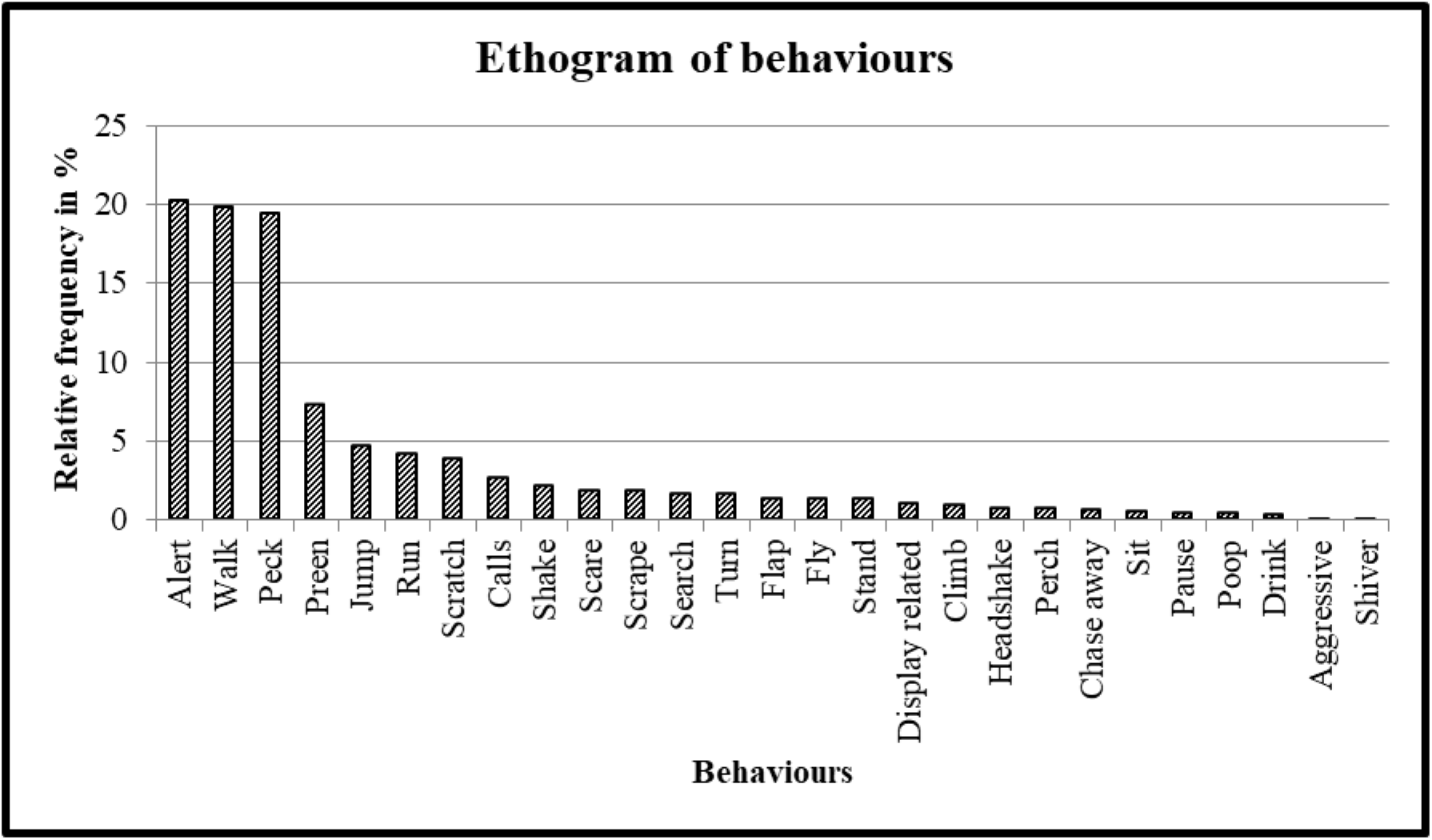
Ethogram of behaviours of Indian Peafowl across all field sites.

For calculating benefit: cost ratio, time spent in feeding was considered as “benefit” while time spent in all other activities (except standing inactive) was considered as “cost”.

### Statistical analysis

For statistical analysis, STATISTICA™ software version 13.2 (Dell Inc. 2016) was used. As the data did not follow normal distribution, non-parametric tests were used for further analysis. Kruskal-Wallis ANOVA was used to see the effect of field sites (Morachi Chincholi, Nashik, Rajasthan) on time spent in feeding, time spent in vigilance, walking and other behaviours. Mann-Whitney U test was used to estimate the effect of proximity to food provision site on time spent in feeding, vigilance, walking and other behaviours.

Benefit: cost ratio was compared across three field sites (Morachi Chincholi, Nashik, Rajasthan) and between food provision and non-provision sites using non-parametric tests such as Kruskal-Wallis ANOVA.

## Results

### Perceptions about Indian peafowl and their relation to agriculture

The age of survey respondents varied from 15-65yrs, all age classes in this range were represented almost equally. Respondents included 90.73% males and 9.27% females. Sources of income for the respondents included agriculture (51.32% of respondents), animal herding (17.88%), labour (farm or other labour 20.2%) and other sources (10. 6%) such as contractor, company job, mechanic, shop owner, artisan, teacher etc. A majority of farmers in the area around RTR are small farmers with 84.11% farmers owning less than 5 acres of land, while rest of the farmers (15.89%) owned 5+ acres to 20+ acres of land. Farmers take two to sixteen types of crops in a year depending on size of farm, availability of labour, water, etc. Most common crops in this region include wheat, bajra, chilly, mustard, sorghum, sesame. All farmers in this region reported crop losses irrespective of time of year during which survey was conducted. The reasons for crop loss differed according to season during which interviews were conducted. Table 3 gives details of season-wise reasons of crop loss. Major crop pests reported in the survey are Wild boar (*Sus scrofa*), Nilgai/ Blue bull (*Boselaphus tragocamelus*), Indian peafowl (*Pavo cristatus*) and Deer. However, number of respondents reporting particular types of crop pests changed according to season in which survey was conducted (Fig 2). Although Indian peafowl is the third common crop pest reported, only 3 % of respondents in November 2016 survey said it was a crop pest compared to 15 % respondents in March 2017 and 20% respondents in July 2017. On the other hand wild boar and blue-bull were reported as crop pests throughout the year by 19-28% respondents. The range reflects variation in data collected across different seasons in the study area.

**Table 3:** Typical reasons of crop loss in RTR region of Rajasthan

| Typical reasons of crop -loss | July-August (%) | March (%) | November (%) |
| --- | --- | --- | --- |
| Crop pests | 62.34 | 59.82 | 100.00 |
| Untimely rains | 36.36 | 31.25 | 0.00 |
| Disease | 1.30 | 8.93 | 0.00 |

**Fig2.**
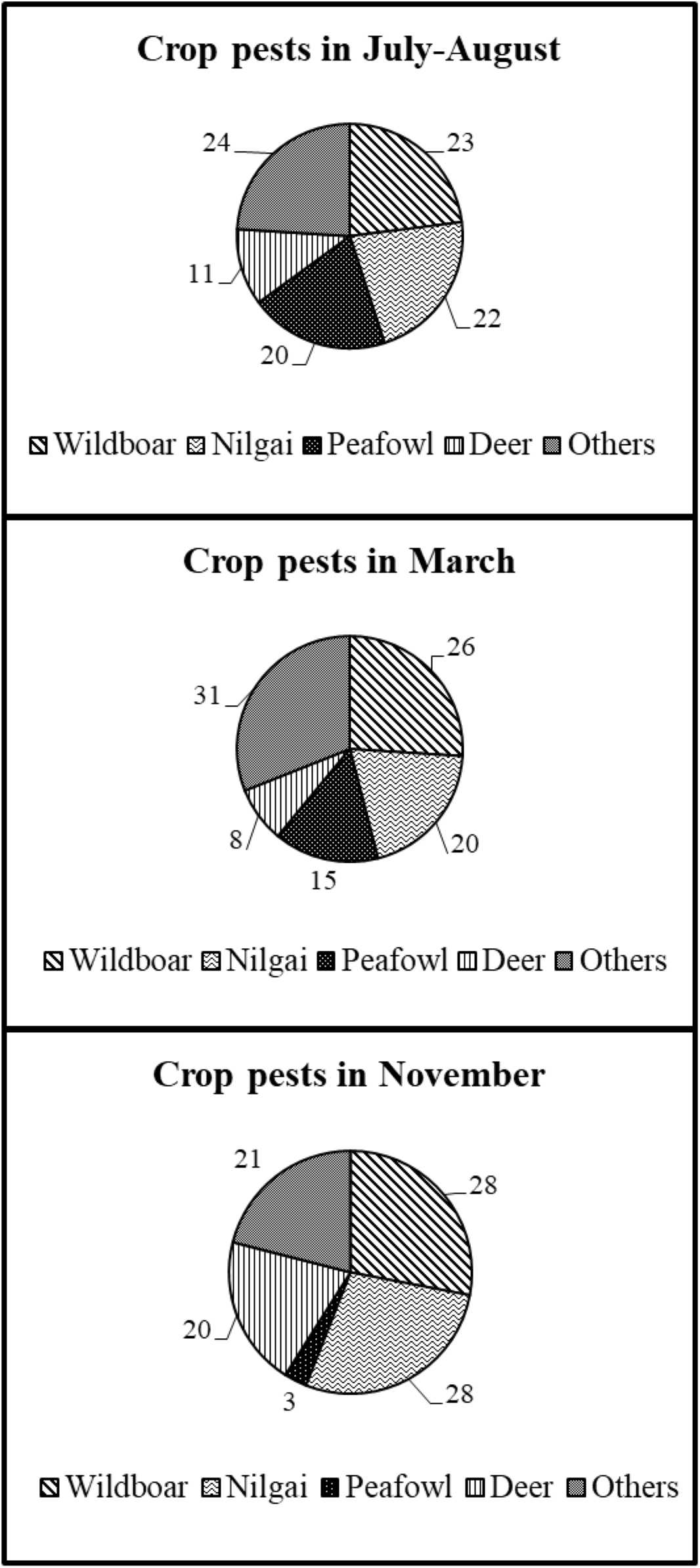
Percentage of respondents reporting crop pests during the months of November, July-August and March in Rajasthan

According to the respondents, peafowl used the agricultural land for feeding and as a source of water (Table 4). Some of the respondents (0.33%) also mentioned that they spread diseases. Between 67.65-100% respondents reported that the long train of feathers causes some damage to crops when the males with full grown train walk through the standing crops. Estimates of total loss due to peafowl varied between 0% to more than 20% according to season in which the survey was conducted (Fig 3). Chickpea, chilli, wheat and mustard were the major crops that incurred loss due to peafowl. Loss was also reported occasionally for coriander, sorghum, sesame, tomatoes, green peas and moong (all reported by 10% or less respondents). Indian peafowl ate crops at post growth (around harvest) stage according to 60-95% of respondents, next most vulnerable stage was pre-growth stage (freshly sown, sprouted or sapling stage) according to 3-29 % of responses. Damage to crops was reported least in the growth stage (8 to 10 % respondents). Very few respondents reported damage to crops at all stages (2 %) and no stages (2 %). (Fig 4). Between 64.65%-100% farmers reported that they changed crop pattern to reduce the loss due to peafowl. Many farmers chose to take a different variety of crop that is less preferred by peafowl (e.g. hybrid chilli instead of indigenous variety) or avoided taking certain crops (e.g. chickpea, ground nut, chilli) or change the place where they take the crop if they own patches of farm lands in more than one locations within the village. Interestingly, the estimate of loss (estimated % loss) due to Indian peafowl showed positive correlation with size of farm (Spearman rank order correlation *r* = 0.24, *p* < 0.05). Farmers with more farm land estimated more crop losses due to peafowl compared to farmers who owned less land.

**Table 4:** Agricultural land usage by Indian Peafowl in RTR region of Rajasthan

| Peafowl's Activities in farm | % of Respondents |
| --- | --- |
| Eat harvest | 42.67 |
| Eat leaves | 17.00 |
| Eat insects | 19.67 |
| Drink water | 6.67 |
| Eat flowers | 8.00 |
| Eat seeds | 0.67 |
| Eat sapplings | 3.67 |
| Nothing | 1.33 |
| Spread disease | 0.33 |

**Fig3.**
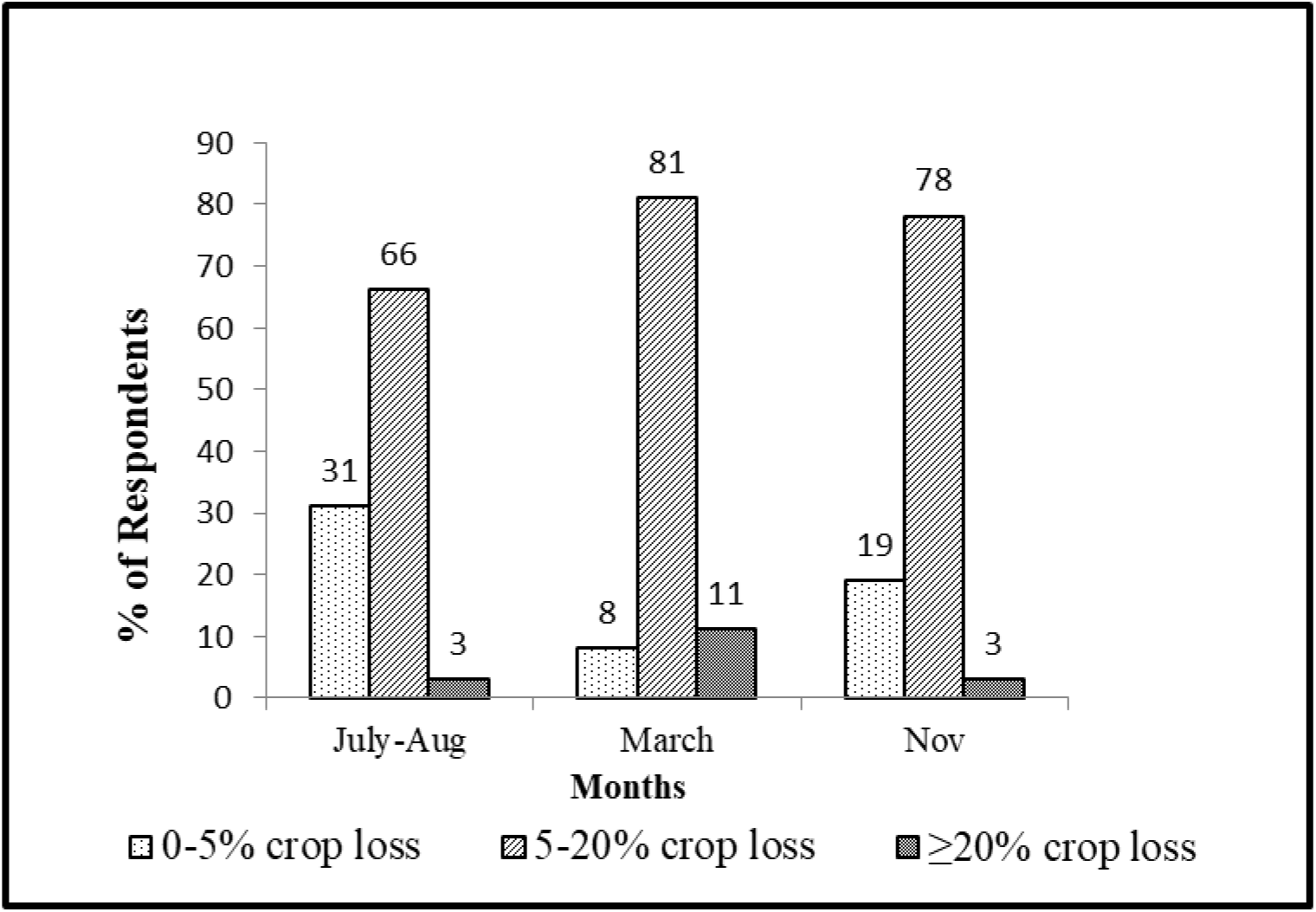
Percentage of respondents reporting crop loss (in %) due to Peafowl during the months July-August, March and November in Rajasthan.

**Fig4.**
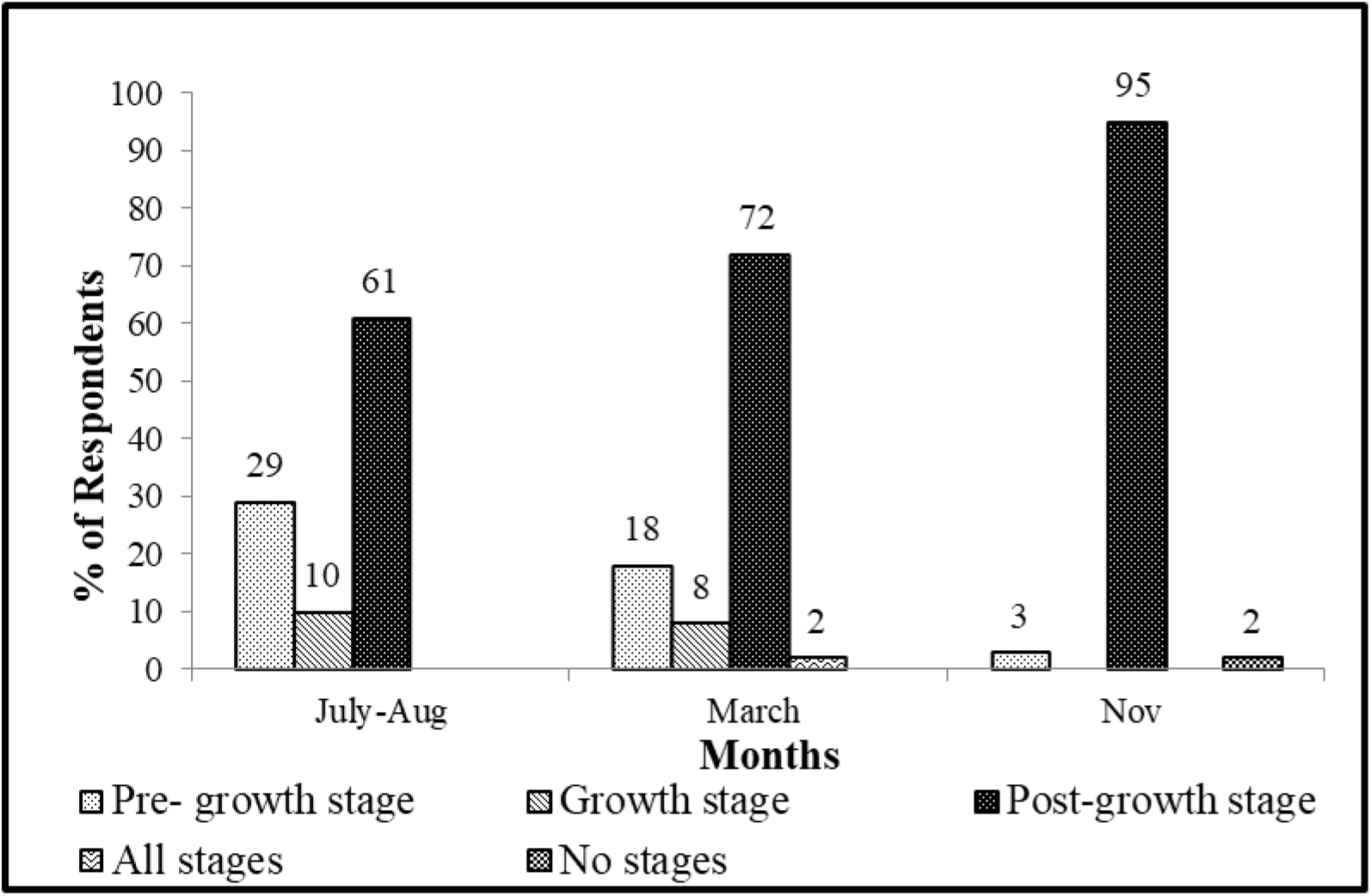
Percentage of Respondents reporting stage of crop affected due to Peafowl during the months July-August, March and November in Rajasthan

### Food provisioning for Indian peafowl

In spite of the crop damage the peafowl cause, local people offer grains to peafowl throughout the year. All households (100% respondents) in the village offer grains to peafowl except in the month of November slightly less number of respondents (94.87%) reported giving grains for peafowl. Most common grains offered to peafowl include bajra (35-45% respondents), sorghum (27.5-41.25% respondents), wheat (10-25.95% respondents), and lentils (0-7.57% respondents) while some people also offer rice, maize, sesame when available. People typically offer a handful (100 – 130 g depending on type of grains offered) to up to 3 Kg grains at a time. In some villages, grains are offered from each household every day, in some cases they are offered in temple once or twice a week or on special days/ occasions. So year around there are on average ~15 Kg grains available for wild life every day in and around temple premises in RTR region.

### Impact of food provisioning on peafowl populations

Impact of food provisioning was studied in three study areas-Morachi Chincholi, Nashik and Rajasthan (RTR). Diet composition of the peafowl was different in the selected study areas. In Morachi Chincholi, peafowl diet consisted of 71% food provisioned by humans compared to 61% provisioned food in Rajasthan, while just 40% peafowl diet in Nashik consisted of provisioned food (Fig 5). Peafowl in Nashik ate almost double the amount of natural (non-grain) food items (60%) compared to peafowl in Morachi Chincholi (29%). Major portion of food provided to peafowl populations in Morachi Chincholi and Rajasthan consisted of grains (66% and 58%, respectively). Peafowl in Rajasthan were not seen eating processed food as opposed to peafowl in Morachi Chincholi and Nashik study area (Table 5).

**Fig5.**
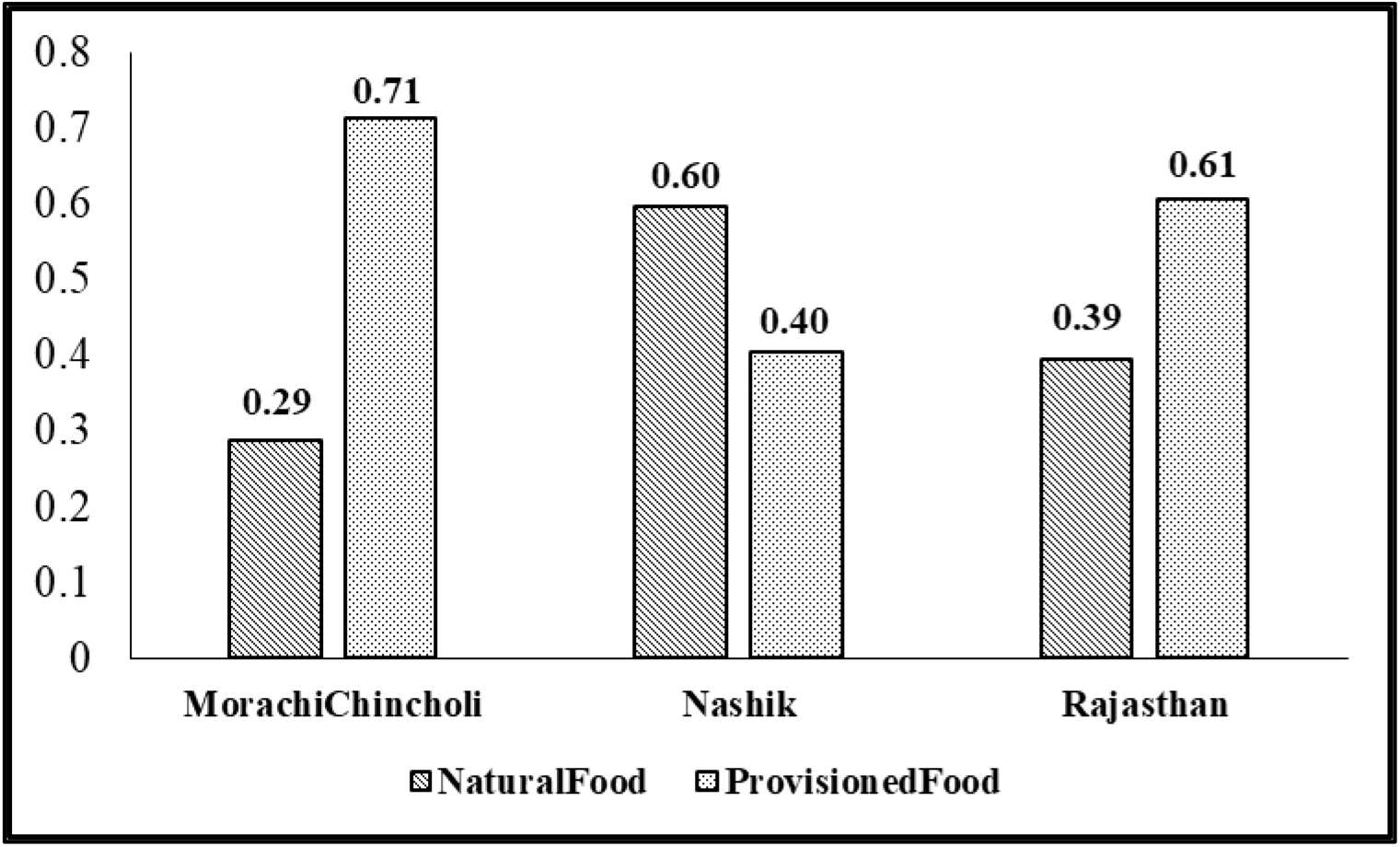
Proportion of natural versus provisioned food items in the Indian Peafowl diet across study areas.

**Table 5:** Diet composition (in %) of Indian Peafowl across field sites

| Field Site | Cereals (%)<br>(provisioned) | Pulses (%)<br>(provisioned) | Processed (%)<br>(provisioned) | Non-grain (%)<br>(non-provisioned) |
| --- | --- | --- | --- | --- |
| Morachi<br>Chincholi | 66 | 3 | 1 | 30 |
| Nashik | 38 | 0 | 2 | 60 |
| Rajasthan | 58 | 3 | 0 | 39 |

There was lot of variation in the total duration of observations for each individual (Range: 1 min-29 min.), however, total observation time did not differ significantly across field sites (Median Test, *Chi-Square* = 1.75, *df* = 2, *p* = 0.42) which confirms that equal efforts were put into observations at the selected field sites. The relative proportion of time spent in different behaviours at food provision sites (where humans put grains for peafowl) and away from provision site (non-provision sites) is shown in Figure 6. Time spent in vigilance (21.88%) and other behaviours (8.98%) at provision sites was comparable to time spent in those behaviours at non-provisioning sites (30.43 %, and 14.26 %, respectively) (Fig. 6). At the provision sites (N=181), individuals spent significantly less time in walking (18.41%) compared to non-provision sites (N=99, 22.99%, Mann-Whitney U test, *p*= 0.003) (Fig. 6 and Fig. 7). Peafowls spent significantly more time in feeding (50.73%) at the provision sites (N=183) as compared to non-provision sites (N=92, 32.32%, Mann-Whitney U test, *p*= 0.0003) (Fig. 6 and fig. 7). Time spent in other behaviours was significantly more at Nashik (N=59) as compared to Morachi Chincholi (N=139) and Rajasthan (N=88), respectively (Kruskal-Wallis ANOVA, Multiple comparison of means, p= 0.03).

**Fig6.**
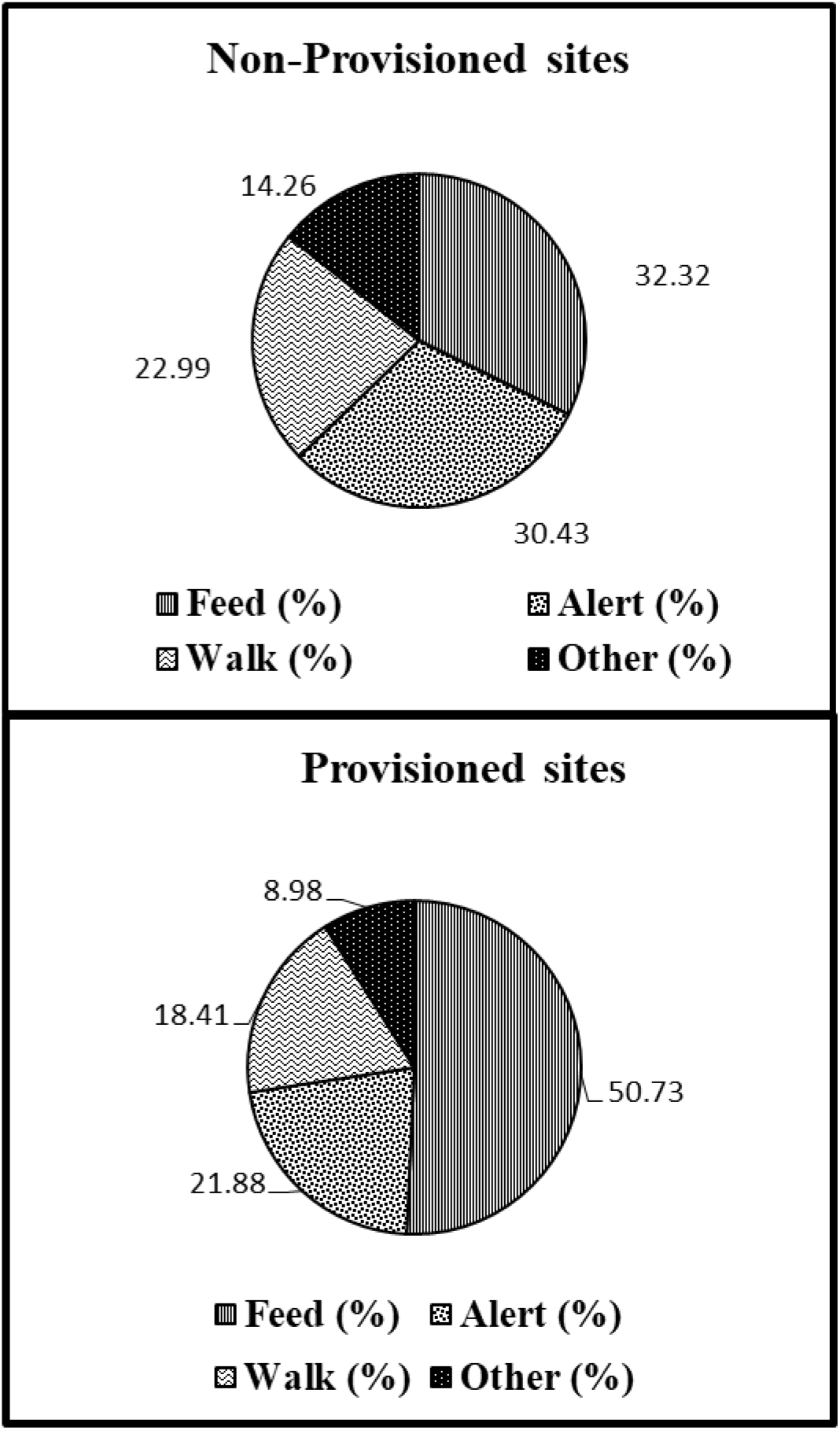
Time budget of Indian Peafowl populations at food provision and non-provision sites.

**Fig7.**
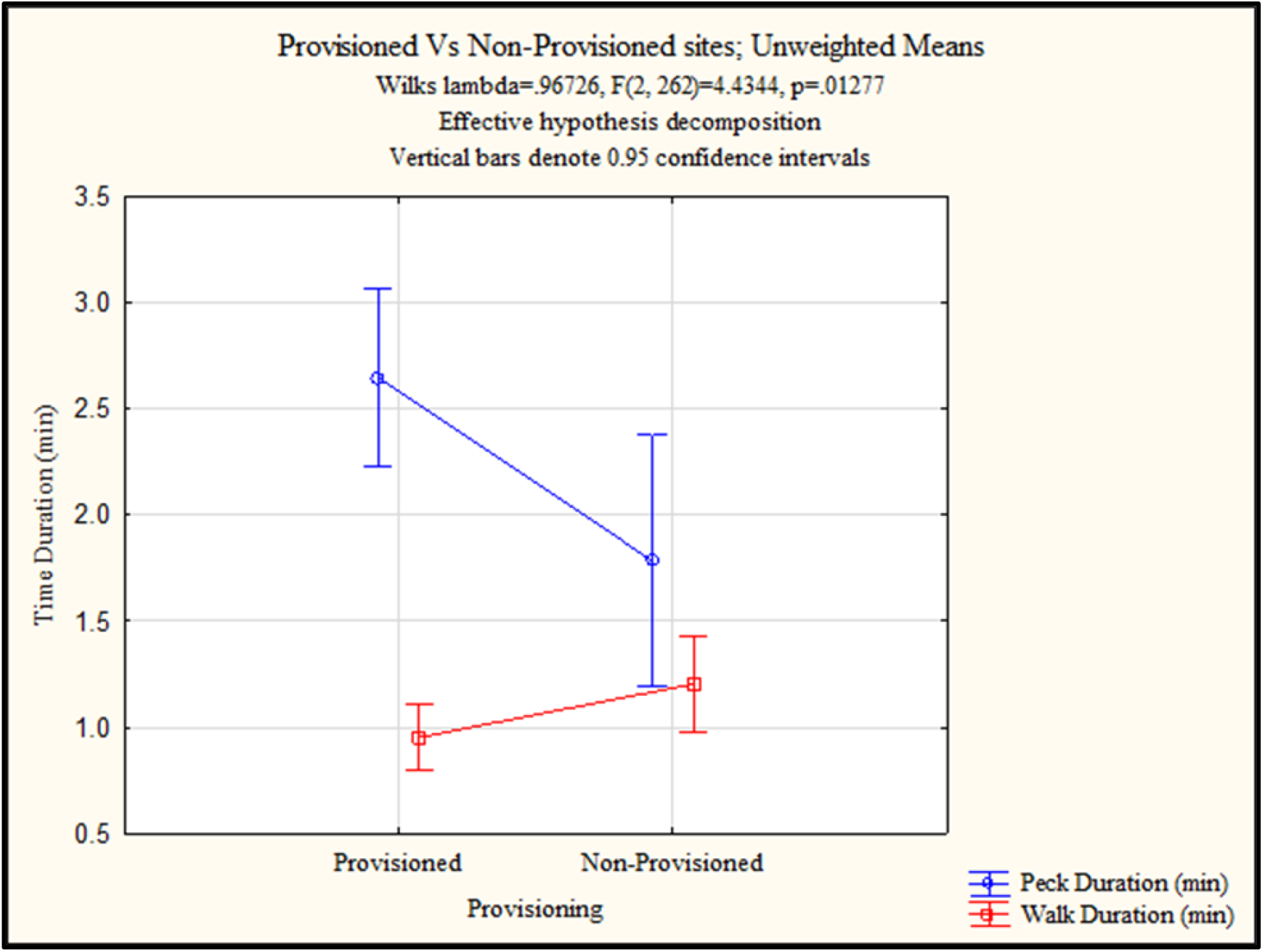
Comparison of time spent in pecking and walking at food provision and non-provision sites. Error bars indicate 95% confidence intervals.

Benefit: cost ratio measured in terms of time spent in feeding (benefit) versus other behaviours (cost) was significantly different between food provision and non-provision sites (Mann-Whitney U test, *p* = 0.00015). The ratio was similar across field sites (Kruskal-Wallis ANOVA, *p* = 0.22).

## Discussion

Human-wild life interactions have mostly been studied so far in the context of human-animal conflict which has drastic consequences such as loss of property, crops or even death. All human-wild life interactions do not necessarily lead to obvious conflicts, nonetheless, they might impact both sides in significant ways. Peoples’ perception about wild life can potentially influence the long-term outcome of such interactions. Our study investigated in detail some aspects of non-violent human-wildlife interactions by focusing on Indian peafowl populations living in human dominated landscape. Our survey covered the perceptions about Indian Peafowl in a wide range of age groups (15-65 yrs) in a geographic area with high population density of peafowls. Most of the surveys were conducted in remote rural areas of Rajasthan by volunteers going door-to-door for collecting data. Survey was conducted considering family as a unit respondent. In most cases, head of the family along with one or two other members (male/ female) present in the household answered the questions. Number of female respondents in our survey remained below 10%. So the results may be biased for male gender. Most of the respondents in the survey were small farmers with less than five acres of land. The source(s) of income, in most cases (89.4% respondents) were related to agriculture (crops + farm animals + labour).

### Perceptions about Indian peafowl and their relation to agriculture

We did not want to bias the respondents by directly asking them if peafowl damaged their crops. Hence, we asked questions step-by-step about types of crops and typical reasons for crop loss in RTR area. Losses due to erratic rain patterns (deficient rainfall, wet drought, untimely rains) is the main reason for crop losses for Indian agriculture which depends largely on Monsoon rains from Indian ocean. Surprisingly, damage due to crop pests emerged as the top most reason across all seasons for loss to farmers in RTR area, followed by loss due to untimely rains, crop disease, or any other reasons (Table 3). The area in which we conducted surveys includes buffer zone and villages just outside the Ranthambhore National park. As a result, wild herbivores from the protected area frequently enter the villages, farmlands and surrounding areas. This can create serious problems for the farmers whose livelihood depends on agricultural produce. Similar issues of crop raiding by wild herbivores have been reported all over the world (Naughton-Treves 1997; Treves et al. 2006; Graham et al. 2010; Mathur et al. 2015; Perera and Vandercone 2017) including India (Chauhan and Singh, 1990; Chauhan et al. 2009; Ramkumar et al. 2014; Meena 2017). Wild boar (*Sus scrofa*), Nilgai/ blue bull (*Boselaphus tragocamelus*), Indian peafowl (*Pavo cristatus*) and deer were reported as the major crop pests in RTR area. While wild boar and blue bull are some of the commonly known crop raiders in India, ours is first systematic survey in which Indian peafowl has been reported as a crop pest in India. There are sporadic newspaper reports and anecdotal information from different parts of India about Indian peafowl damaging crops (Ganesan, R. 2016), however, there has been no systematic survey or documentation of extent and type of crop damage done by Indian peafowl as we have done in this survey. Recently, Indian peafowl was reported as a crop pest by farmers in Sri Lanka (Horgan et al. 2017; Perera and Vandercone 2017). One study reported that Maavee paddies and surrounding habitats offer refuges for wildlife including Indian peafowl; and Indian peafowl damaged rice at the seedling and grain filling stage irrespective of the farming method used (Horgan et al. 2017).

Peafowl used agricultural land as a source of food, water (Table 4), display sites and resting sites (personal observations). A majority of respondents (87.63-95.45%) in our survey reported that peafowl do affect crops in an adverse way. So Indian peafowl is not only visiting their farms seeking refuge or using farms as new feeding habitat, in most cases they do have perceivable negative effects on crops. Although they were not reported to spread disease (only 0.33% respondents think they spread crop disease), their long train was found to cause some damage as they walked through the standing crop. They also eat various parts and stages of the crop (Fig. 4). Estimates of total crop loss due to Indian peafowl ranged from negligible (0%) to over 20%. Typically, figures between 5-20% losses were mentioned by respondents for various crops. It is important to note that these are “perceived” estimates of loss given by farmers and the actual figures need to be verified through more detailed ‘on field’ data collection. Other wild herbivores such as elephants can cause up to 35% crop damage (Ramakumar et al. 2014), while wild boars can destroy 5-36% of crops (Chauhan et al. 2009). Blue bull (*Boselaphus tragocamelus*) seems to have the highest impact with 58% losses (rarely less than 10%) in Harayana, India (Chauhan and Singh 1990). The losses due to blue bull can be as high as 70% in high density areas (Meena 2017). Comparatively, 5-20% loss due to Indian peafowl seems less, nevertheless, it may still be considerable amount of loss to small farmers in RTR area. Chickpea, chilli, wheat, mustard (17-50% respondents for each crop) are the crops preferred by Indian peafowl compared to tomatoes, sesame, coriander, sorghum, green peas, moong (less than 10% respondents).

Peoples’ perception about Indian peafowl being a crop pest changed according to season in which they were interviewed. While wild boar or blue bull were consistently reported as crop pests throughout the year, only 3 % respondents mentioned Indian peafowl as crop pest in November, 15% in March and as many as 20 % respondents called it a crop pest in July 2017. Our survey also indicates that crops around harvest stage (post-growth stage) are most likely to be eaten by Indian peafowl followed by pre-growth stage (freshly sown, sprouted or sapling stage) and growth stage (between saplings to leaves set). Least damage is incurred due to peafowl visiting the growing crops (Fig 4).

The crop damage at different stages of growth may lead to different amount of loss. Some of the damaged parts may regenerate but there is nevertheless a substantial amount of loss in terms of total yield (Bayani et al. 2016). In contrast, for some crops such as chickpea, herbivory in first 20 days after sowing may actually lead to greater branching resulting in more number of seeds. However, herbivory beyond a threshold in pre-flowering stage is known to decrease crop yield even in chickpea by as much as 67% (Bayani et al. 2016).

Many farmers have adopted different strategies to reduce the crop loss due to peafowl. The strategies include guarding the farms during most vulnerable stages, changing crop variety or place. The effectiveness of these strategies needs to be checked thoroughly. None-the-less the strategies to reduce crop damage/ loss themselves cost the farmers in terms of time, resources, money invested not to mention the inconvenience and loss incurred due to choosing less than optimal choice of crop or farming place.

Overall, the effects of crop damage due to Indian peafowl may vary with type of crop taken, season, etc. There is no denying that Indian peafowl seems to be a crop pest at least in some parts of India. Surprisingly, most respondents in our survey did not have negative views or resentment about Indian peafowl damaging/ eating their crops. Many of them, in fact, mentioned that even peafowl have “right-to-live” and farmers do not consider it as “loss” if peafowl come and eat in their farm. It is possible that the ‘perceived beauty’ of peafowl along with their association with popular deities in India (Nair, 1974) and relatively less amounts of crop damage by this species may play a role in biasing the perception of people towards this species as compared to the other pests. This seemingly positive perception is in sharp contrast to farmers’ perceptions when other crop pests such as wild boar, elephants or blue bulls visit their crops. Most of the households in RTR area in fact pro-actively offer grains to peafowl throughout the year (94.87-100% respondents). Typically bajra, sorghum, wheat or lentils are offered by each household in a designated place around temple premises in RTR region. The quantity or frequency with which each household offers grains may vary from place to place but on average about 15 Kg of grains are offered daily at these designated places in RTR region as part of temple ritual/ traditional belief. As a result, such places have become a reliable source of food for the wildlife including Indian peafowl. Thus, people’s positive/ neutral perception about Indian peafowl, traditional beliefs and ritual practices such as active food provisioning at temples may be important for maintaining high population numbers of peafowl in RTR area. One can conclude that human-wildlife interactions and people’s perception/ attitude towards wild life can impact not only food provisioning but also wild life management (Jacobs et al. 2014a, b), conservation efforts (Jacob and Harms 2014) and ultimately human–wildlife relationships (Jacobs et al. 2012).

### Effects of food provisioning on peafowl populations

All around the world humans have provided many novel and reliable food resources to herbivores in the form of crops and ornamental plants. Some species exploit these resources quite well and can become economically important crop pests (Sih et al. 2011). Generalist herbivores (those who have broad range of plant species they like to feed on) are more likely to exploit the novel food resources (crops, ornamental plants) present in the human dominated landscapes. Peafowls generally have a broad range of diet options including plant parts (fresh sprouts, leaves, flowers, fruits, seeds) as well as insects, worms, sometimes small reptiles (Sathyakumar and Kaul 2007). In spite of having most of these diet options available in our study areas, peafowls were found eating food provided by humans (cereals, pulses, processed food). According to our estimate, ~30-35 Kg food grains are offered to peafowl per day in the village and surrounding areas of Morachi Chincholi, while ~15Kg grains are offered per day around homes and temple premises in the villages in Rajasthan included in this study. In both these study areas, grains are offered at designated places everyday throughout the year. Thus, Indian peafowl population in Morachi Chincholi has access to diet rich in carbohydrates (cereals) and proteins (pulses) throughout the year (Table 2). As much as 71% of their diet consisted of food provisioned indirectly in the form of crops or directly as grains offered in the village surroundings (Fig. 5). Villages in Rajasthan offer relatively less variety of cereals and very few pulses, yet the diet of peafowl in Rajasthan has up to 61% of food provisioned by humans versus 39% natural food. Peafowl population in Nashik, on the other hand has access to less reliable and less varied food offered by humans (~ 3-5 Kg per day), as a result only about 40% of their diet consists of provisioned food. It is argued that crop varieties are rich in nutritional quality and poorer in secondary metabolites compared to their wild counterparts (Rode et al., 2006). If this is true, it could explain why peafowl are choosing harvested/ unharvested crops and ready to eat grains/ food, wherever they are available, over their “natural food”. The quantity, variety and nutritional quality of food available to peafowl plus the reliability of such food resources might be crucial factor in making their food choices and overall diet composition (Table 5).

Food provisioning not only influenced their diet composition, but also the time budget. At food provision sites, the peafowls could spend up to 50.73% of their time just eating, which is significantly higher than the time spent in feeding at non-provision sites (Fig. 6 and fig. 7). They could get high quality food in one place without spending much time in walking. In contrast, the peafowl had to spend more time in walking in search of food if they were feeding at non-provision sites, while they could spend only 32.32% of their time in feeding (Fig. 6). Effect of less amount of food provisioning at Nashik as compared to Morachi Chincholi and Rajasthan can be seen in the higher time spent by Peafowls in other behaviours. Our results suggest that the benefit: cost ratio in terms of time spent in behaviours can be influenced due to food provisioning. We could not calculate benefits and costs in terms of energy budget as it was difficult to estimate energy gain from non-provisioned food items. It will be interesting to explore other cost benefits of food provisioning in terms of body condition/ health of Indian peafowl near human habitation.

Effects of food provisioning on time budgets, diet and activity seen in Indian peafowl populations are very similar to effects seen in primate populations (Kamal et al. 1997; Berman and Li 2002; Ram et al. 2003; Hadi et al. 2007; McKinney 2010; El-Alami et al. 2012). Interestingly, these effects extended to the ranging patterns, foraging success, and consequently fitness of primates (Oro et al. 2013; Becker and Hall 2014). Food provisioned primate troops also had higher growth rates and population densities (Altmann and Muruthi 1988; Jaman and Huffman 2013). Comparable studies on birds, however, are rare especially in Indian subcontinent. Thus, our study for the first time has documented effects of food provisioning on feeding ecology of big bird such as Indian peafowl.

Supplementary food available at bird feeders (typically in western countries) is known to have far reaching consequences on avian ecology in terms of increasing survival during overwintering, enhanced breeding success, changing sex-ratios of offspring in smaller avian species, range expansion of species (Robb et al. 2008 and references therein). Although, most of these consequences are positive, some negative consequences of food provisioning are also possible in avian as well as primate systems. If animals start depending on food provided by humans it may lead to behavioural changes as illustrated in Finland where some great-tit populations are so dependent on supplementary food during winter that they can no longer be sustained by natural food resources alone (Orell 1989). In Morachi Chincholi, where large amounts of grains are provisioned throughout the year, somewhat parallel scenario is seen during summer months. In the months of April, May and part of June, there are high temperatures (up to 42-45°C), water is scarce and if there are no crops during summer months in a particular year, the peafowl concentrate around remaining waterbodies (village wells, ponds) and wherever humans provide them grains and water (personal observations). Although it is not surprising that peafowl incorporated nutrient rich resources in their diet wherever such option was available, it is interesting that these resources are being actively provided to them by humans in at least some places. It may, thus, be important to factor in human attitude towards this species in population management or future conservation efforts especially considering how frequently they are found in human dominated landscape.

The dynamic interplay of experiential, ecological, socioeconomic and cultural factors can influence perception and local human behaviour towards wild-species with significant implications for the management of wild species and the ecosystems they inhabit (D’Lima et al., 2013). Studies such as ours may help in designing effective conservation strategies for a magnificent bird that intrigued even Charles Darwin.

## Supporting information

Supplementary Questionnaire

## Acknowledgements

Thanks to Maharashtra Education Society’s Abasaheb Garware College, Pune for hosting the Ramalingaswami Re-entry Fellowship to DAP and providing the infrastructure and administrative support. DAP and PD are thankful to Late Mr. Biswaroop Raha for suggesting field sites in Nashik, Dr. Dharmendra Khandal and volunteers of Tiger Watch who extended support for field work in Rajasthan. Authors also appreciate the support and hospitality of Morachi Chincholi residents during field work. We are thankful to Rupesh Gawade, Pranav Mhaisalkar, Ankita Divekar, Akash Dubey and Vedanti Mahimkar for helping with data collection on field.

## Declarations

### Authors contributions

DAP conceived the study, designed methodology, collected data, analysed the data and prepared the manuscript. PD collected data, helped with analysis and manuscript preparation.

### Permissions

No live animals/ samples were handled during this study. The study was conducted outside protected areas. Therefore, the work did not require special permissions.

