## Supplementary Questionnaire for "A tale of two species: human and peafowl interactions in human dominated landscape influence each other’s behaviour"

**Economic Impact Survey Questionnaire**

Date of Survey:

**Introduction protocol:**
- Good morning/ afternoon/ evening/ Ram Ram
- I am xxxxxx doing research work at xxxxxx. As part of my research work, I am studying impact of peacocks on the local population and economy. This study will help us understand dependence of humans and animals on each other.
- If you could spare 15 min. to give me relevant information it would be of great help for the study.
- I would like to assure you that this information is being gathered purely for study/ research purpose and it will be treated in the strictest of confidence.

**Oral Consent:**
Would you like to participate in the study right now? Or at other time convenient to you?

I. **Basic information:**
   1. Name:
   2. Age:
   3. Sex:
   4. Address:
   5. No. of family members:

II. **Livelihood:**

1. How many family members are there?

2. What is the source(s) of income? (please tick mark all that apply)
   Farmer (owner)
   Farm worker
   Business (details)
   Job (details)
   Other (details)

3. What is the yearly income from all the sources?
   Less than Rs.100000
   Between Rs.100000-Rs.500000
   More than Rs.500000

**For farmers go to question 4, for others, go to part III. (Mark all that applies)**

4. What is the size of farm?

5. Which crops have you taken through-out the last year?

Grains
Cereals
Fruits
Veggies
Sugarcane
Others (please specify)

6. Do you grow crops for household or do you sell it?

7. How much is the expected yield for each crop approximately?

8. Are there crop losses? (YES/NO)

9. If yes, what are the typical reasons of crop loss?
   Drought/ water scarcity
   Crop disease
   Crop pests
   Untimely rain
   Shortage of farm labor
   Other reasons (please specify)

10. If due to crop pests, what type of damage is seen?
    Trampling
    Eating crop parts such as tender leaves, flowers, fruits, roots
    Spreading disease
    Any other (please specify)

11. What type of crop pests do you encounter?
    Wild Pig/ deer/ birds/ cattle/ sheep/rodents/ hare/ any other (please specify)

12. Do peafowl affect crops? (YES/NO)

13. What do the peafowl do on the farm?
    Drink water
    Eat fresh growth/ fruits/ grains/ leaves/ insects/ lizards
    Anything else? (Please specify)

14. Are crops destroyed because of their weight/ big train?
    Yes/ No/ Sometimes

15. Do the peafowl eat crop pests such as insects/ mice?
    Yes/ No/ Sometimes

16. Do you put separate food/ grains/ water for peafowl?
    Yes/ No/ Sometimes

17. If yes, how much grains do you put daily/ weekly?

18. If yes to Q17, which grains/ food do you typically give?

19. What would be % of crop loss due to peacocks (vs. due to other reasons)
    0-5%
    5-20%
    More than 20%

20. Do you use pesticides/ insecticides on crop?

21. Which type(s) of crops do the peacocks go to/ raid more frequently?

22. Which stage of the crop that you take is most susceptible to damage by peafowl?
    Pre-growth stage (sowing to sapling)
    Growth stage (sapling to seed/ fruit set)
    Post-growth stage (seed/fruit set to harvest)
    All stages
    No stages

23. Which season do the peafowl visit farm more often?

24. Do you have to change the crop plan due to peafowl?

25. What do you do to keep away peafowl from crops?

26. How many helpers or farm workers do you have on the farm?

**Questions for non-farmers/ part-time farmers:**

27. Do you gather/ sell naturally shed feathers?

28. If yes, at what price?

29. Do you put separate food/ grains/ water for peafowl?
    Yes/ No/ Sometimes

30. If yes, how much grains do you put daily/ weekly?

31. If yes to Q29, which grains/ food do you typically give?

**III Perception:**

32. We are seeing lots of peacocks here. Since when are they here?

33. What do you think about so many peacocks in the surrounding area?

34. Are there any stories/ anecdotes/ rituals/ traditions about peacocks in the village? (please give details)

35. Are the feathers sold in the village?

36. If yes, typically at what price?

37. Any other peacock “products” (feathers/ eggs/ pictures/ souvenirs) sold?

38. Does anyone kill peafowl?

39. What is the procedure if someone finds out about killing/ harming peafowl in the village?
